# Three orphan histidine kinases inhibit *Clostridioides difficile* sporulation

**DOI:** 10.1101/2021.11.18.469199

**Authors:** Adrianne N. Edwards, Daniela Wetzel, Michael A. DiCandia, Shonna M. McBride

**Affiliations:** Department of Microbiology and Immunology, Emory University School of Medicine, Emory Antibiotic Resistance Center, Atlanta, GA, USA

**Author notes:** Corresponding author. Mailing address: Department of Microbiology and Immunology, Emory University School of Medicine, 1510 Clifton Rd, Atlanta, GA 30322.

**Keywords:** *Clostridioides difficile*, sporulation, spore, anaerobe, histidine kinase, Spo0A, phosphotransfer, phosphorylation, phosphorelay, phosphatase

## Abstract

The ability of the anaerobic gastrointestinal pathogen, *Clostridioides difficile*, to survive outside the host relies on the formation of dormant endospores. Spore formation is contingent on the activation of a conserved transcription factor, Spo0A, by phosphorylation. Multiple kinases and phosphatases regulate Spo0A activity in other spore-forming organisms; however, these factors are not well conserved in *C. difficile*. Previously, we discovered that deletion of a conserved phosphotransfer protein, CD1492, increases sporulation, indicating that CD1492 inhibits *C. difficile* spore formation. In this study, we investigate the functions of additional conserved orphan phosphotransfer proteins, CD2492, CD1579, and CD1949 which are hypothesized to regulate Spo0A phosphorylation. Disruption of the conserved phosphotransfer protein, CD2492, also increased sporulation frequency, similarly to the *CD1492* mutant, and in contrast to a previous study. A *CD1492 CD2492* mutant phenocopied the sporulation and gene expression patterns of the single mutants, suggesting that these proteins function in the same genetic pathway to repress sporulation. Deletion of the conserved CD1579 phosphotransfer protein also variably increased sporulation frequency; however, knockdown of *CD1949* expression did not influence sporulation. We provide evidence that CD1492, CD2492 and CD1579 function as phosphatases, as mutation of the conserved histidine residue for phosphate transfer abolished CD2492 function, and expression of the *CD1492* or *CD2492* histidine site-directed mutants or the wild-type *CD1579* allele in a parent strain resulted in a dominant negative hypersporulation phenotype. Altogether, at least three phosphotransfer proteins, CD1492, CD2492 and CD1579 (herein, PtpA, PtpB and PtpC) repress *C. difficile* sporulation initiation by regulating activity of Spo0A.

**IMPORTANCE:** The formation of inactive spores is critical for the long-term survival of the gastrointestinal pathogen *Clostridioides difficile*. The onset of sporulation is controlled by the master regulator of sporulation, Spo0A, which is activated by phosphorylation. Multiple kinases and phosphatases control Spo0A phosphorylation; however, this regulatory pathway is not defined in *C. difficile*. We show that two conserved phosphotransfer proteins, CD1492 (PtpA) and CD2492 (PtpB), function in the same regulatory pathway to repress sporulation by preventing Spo0A phosphorylation. We show that another conserved phosphotransfer protein, CD1579 (PtpC), also represses sporulation, and we eliminate the possibility that a fourth orphan histidine kinase protein, CD1949, impacts *C. difficile* sporulation. These results support the idea that *C. difficile* inhibits sporulation initiation through multiple phosphatases.

## INTRODUCTION

*Clostridioides difficile* undergoes a significant differentiation process to develop dormant endospores, which enable this anaerobic pathogen to survive outside of the mammalian gastrointestinal tract for a prolonged period of time. The environmental cues and regulatory pathways that govern the initiation of sporulation all converge on Spo0A, the master regulator of sporulation (1–3). Spo0A is a conserved transcriptional regulator present in all endospore-forming bacteria and is essential to this process (4). Spo0A activity is controlled by the phosphorylation of an aspartate residue, allowing Spo0A to directly bind to specific target sequences in the promoters under Spo0A regulation (3, 5, 6). Thus, active Spo0A~P drives transcription of sporulation-specific genes whose products are required for entry into the sporulation pathway (7, 8).

In other spore formers, the opposing activities of numerous orphan histidine kinases and phosphatases contribute to Spo0A phosphorylation, presumably in response to environmental stimuli or nutritional cues. In the well-studied soil bacterium, *Bacillus subtilis*, Spo0A is phosphorylated via an expanded two-component signal transduction system (TCS), known as a phosphorelay (9). The *B. subtilis* phosphorelay is comprised of multiple phosphotransfer proteins which transmit a phosphate from one of several sensor histidine kinases through two phosphotransfer proteins to Spo0A. Phosphatases directly dephosphorylate Spo0A or a phosphotransfer protein in the phosphorelay, or inhibit activation and/or autophosphorylation of the sensor histidine kinases. However, many of the key regulatory proteins that control Spo0A activation in *B. subtilis* are absent from the *C. difficile* genome (10–12), supporting the hypothesis that *C. difficile* controls the initiation of sporulation differently than the *Bacillus* sp.

The *C. difficile* genome does not encode orthologs of the *Bacillus* species’ phosphotransfer proteins, suggesting that either unique orphan histidine kinases directly phosphorylate Spo0A or that other proteins transfer the phosphate from the kinases to Spo0A (10). Reinforcing the former hypothesis, orphan histidine kinases promote spore formation and have been shown to directly phosphorylate Spo0A in Clostridia, including in *C. acetobutylicum* and *C. perfringens* (13, 14). In *C. difficile*, however, only the putative sporulation-associated histidine kinase CD1492 has been studied in depth to ascertain its role in *C. difficile* sporulation (15). A *CD1492* mutant exhibited a hypersporulation phenotype, had decreased TcdA production, and was significantly less virulent in the hamster model of *C. difficile* infection (15). Two other annotated sporulation-associated histidine kinases, the membrane-bound CD2492 and soluble CD1579, were briefly characterized in a previous study (16). This study showed that a *CD2492* mutant has a decreased sporulation frequency via microscopy after extended growth in rich medium and provided evidence that CD1579 directly transferred a phosphate to Spo0A *in vitro*. However, the conclusions of this study in regards to the function of either protein were limited.

Here, we further probed the function of CD2492 and CD1579 in *C. difficile* spore formation, as well as asked whether an additional conserved histidine kinase, CD1949, influences sporulation. Our results revealed that a null *CD2492* single mutant and a combined *CD1492 CD2492* mutant exhibited the same high sporulation frequency and increased sporulation-specific gene expression as the *CD1492* mutant, indicating that these proteins function in the same regulatory pathway. A *CD1579* mutant also exhibited high, but variable, sporulation phenotype. We demonstrate that mutating the conserved histidine residues required for phosphate transfer in each of these putative histidine kinases impact *C. difficile* spore formation in various ways, providing evidence that phosphate transfer is important for CD1492, CD2492 and CD1579 function. Finally, we show that CD1949 does not influence *C. difficile* sporulation. Because the functions of CD1492, CD2492 and CD1579 influence Spo0A phosphorylation, but their phenotypes do not support their primary activities as Spo0A kinases, we propose to name the corresponding loci <u>p</u>hosphotransfer <u>p</u>rotein A (*ptpA*; *CD1492*), <u>p</u>hosphotransfer <u>p</u>rotein B (*ptpB*; *CD2492*) and <u>p</u>hosphotransfer <u>p</u>rotein C (*ptpC*; *CD1579*). Together, these three phosphotransfer proteins prevent *C. difficile* sporulation.

## MATERIALS AND METHODS

### Bacterial strains and growth conditions

The bacterial strains and plasmids used for this study are listed in **Table 1**. *Clostridioides difficile* strains were routinely cultured in a 37°C anaerobic chamber (Coy) with an atmosphere of 10% H_2_, 5% CO_2_ and 85% N_2_, as previously described (17), either in BHIS or TY medium pH 7.4. *C. difficile* cultures were supplemented with 2 to 10 μg/ml thiamphenicol if necessary for plasmid maintenance, and overnight cultures included 0.1% taurocholate to promote spore germination and 0.2% fructose to inhibit sporulation, as indicated (18, 19). *Escherichia coli* strains were grown at 37°C in LB with 100 μg/ml ampicillin and/or 20 μg/ml chloramphenicol as indicated, and 50-100 μg/ml kanamycin was used to counterselect against *E. coli* HB101 pRK24 after conjugation with *C. difficile* (20).

**Table 1.**
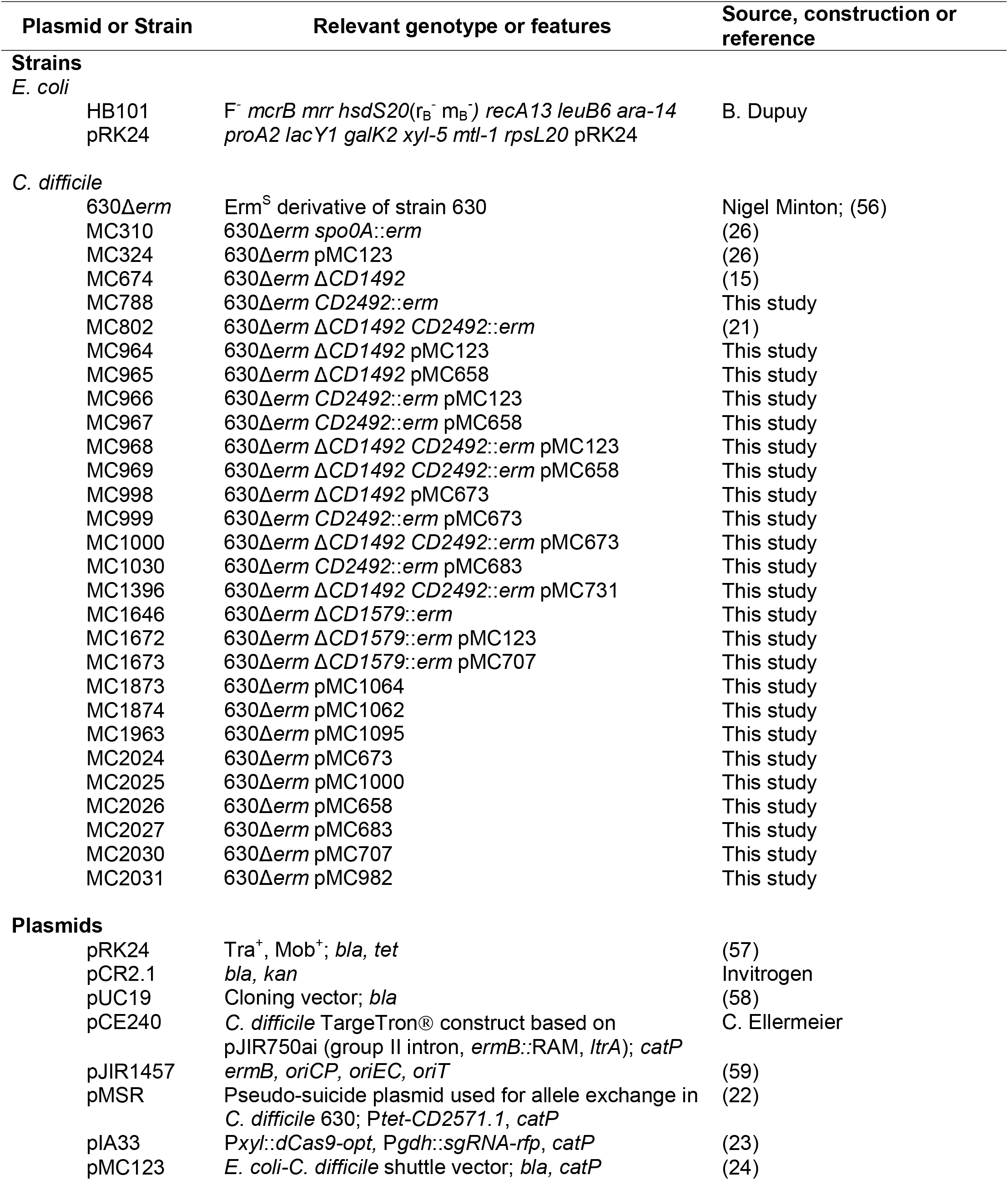
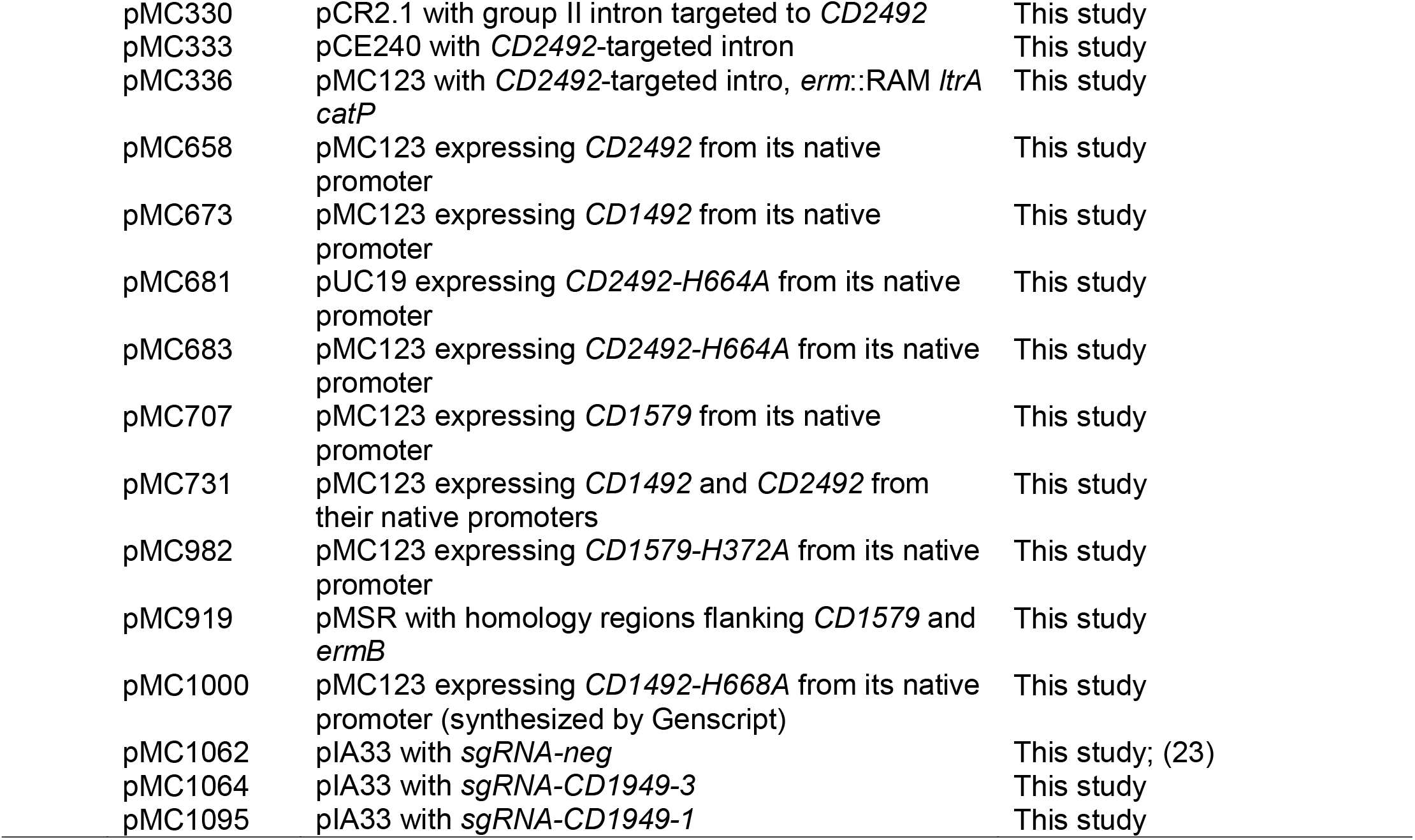
Bacterial Strains and Plasmids.

### Strain and plasmid construction

*C. difficile* 630 (Genbank no. NC_009089.1) was used as the template for primer design, and *C. difficile* 630 was used as the template for PCR amplification and mutant construction. Oligonucleotides used in this study are listed in **Table 2**. The 630Δ*erm CD2492* mutant (*ptpB*; MC788) was recreated by retargeting the group II intron from pCE240 using the targeting site published in Underwood *et al*. 2009 (16). Notably, the targeting site was not located in the 254a site within the *CD2492* coding region noted in Underwood *et al*. 2009, but rather at 318s. The *CD1492 CD2492* double mutant (*ptpA ptpB*; MC802) was constructed similarly using the 630Δ*erm* Δ*CD1492* (*ptpA*; MC674) background (21). All strains were confirmed with PCR analysis (**Fig. S1A**).

**Table 2.**
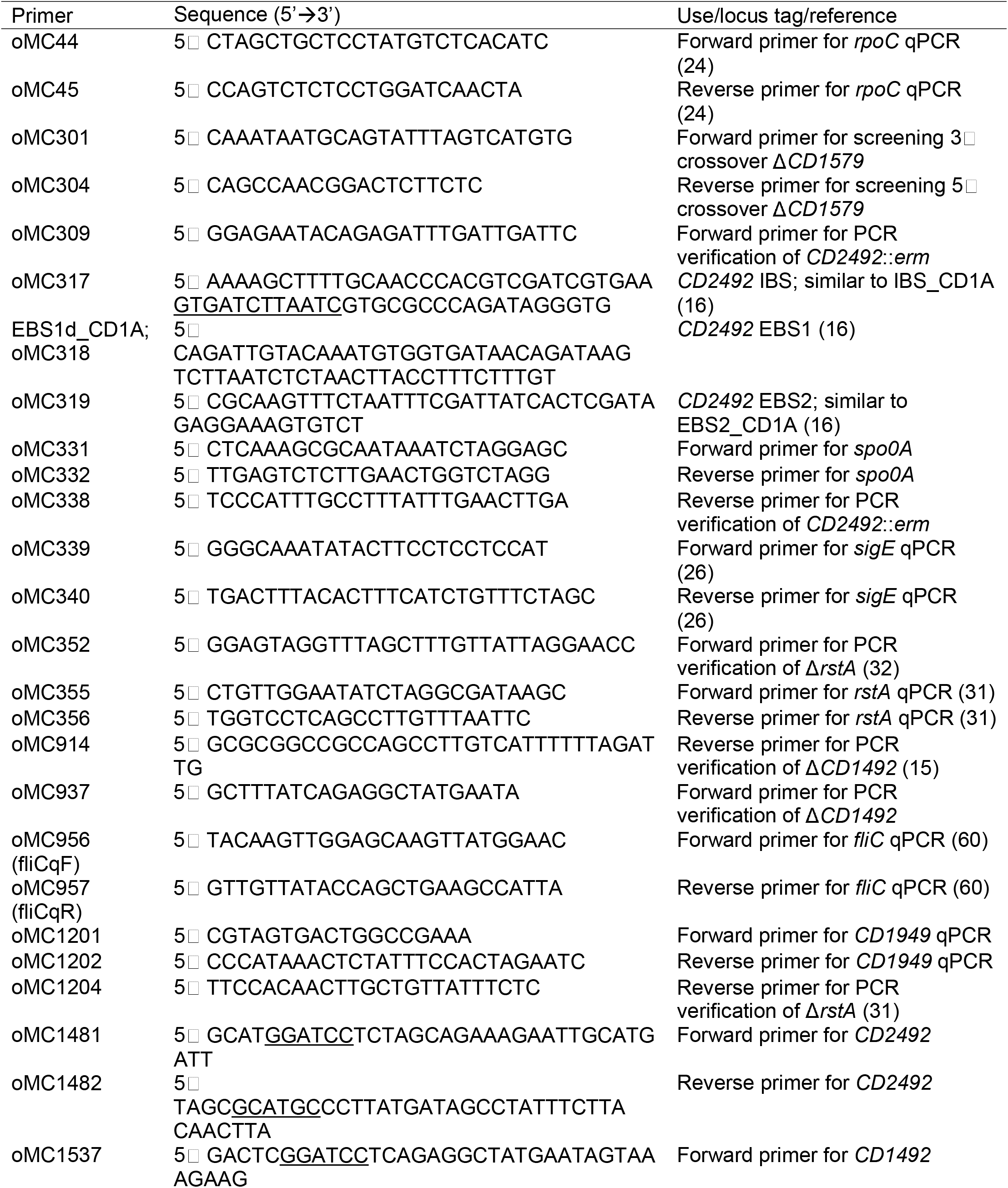
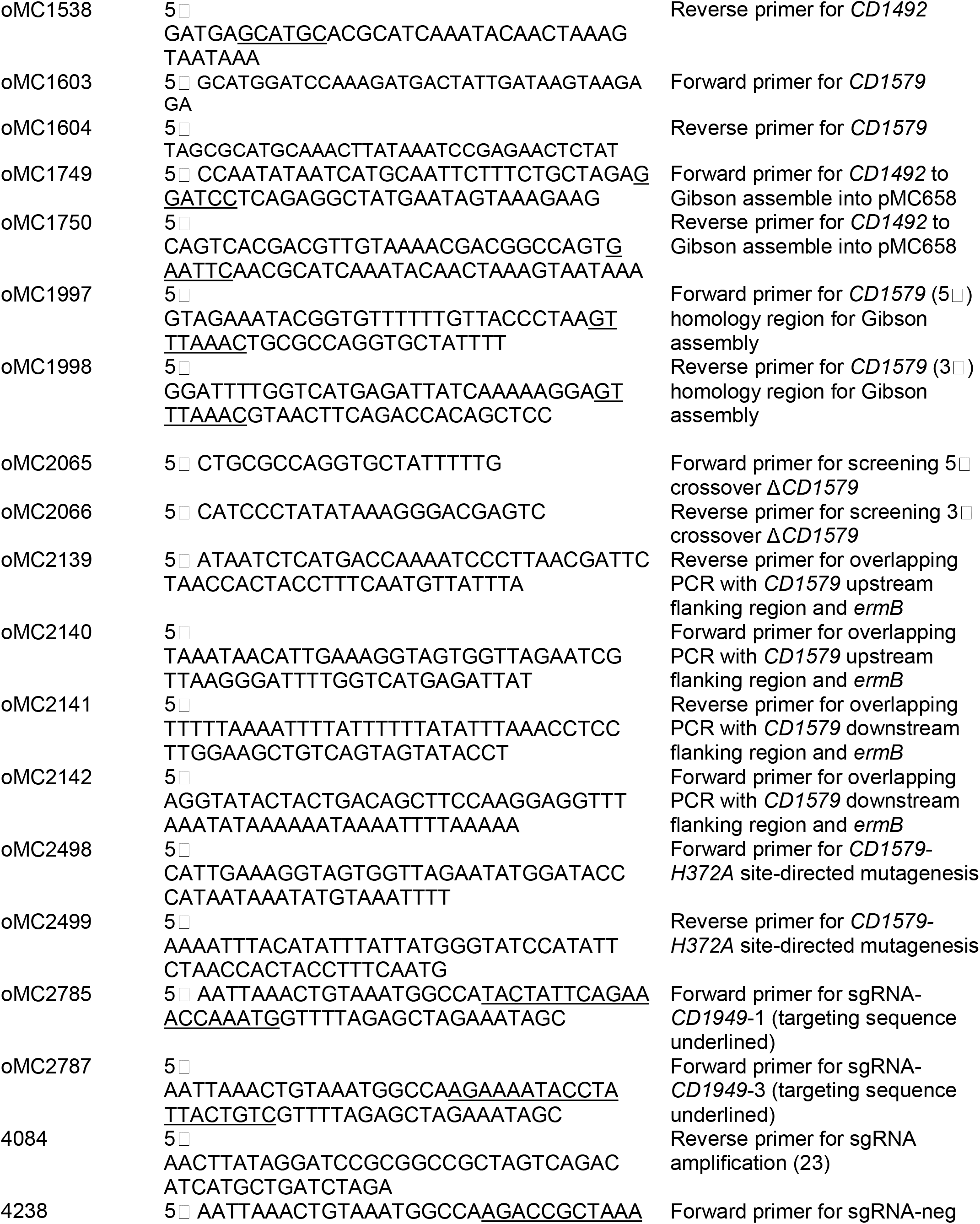
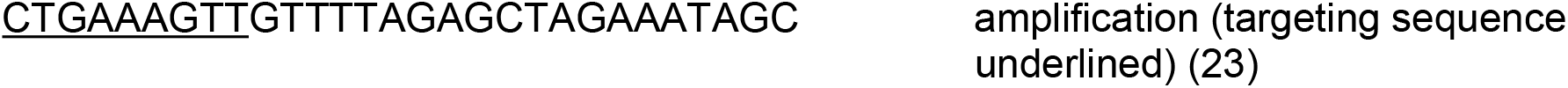
Oligonucleotides.

The *CD1579* (*ptpC*) mutant was created using the pseudo-suicide allele-coupled exchange (ACE) vector as previously described (22), with some modifications. A pMSR-derived vector, pMC919, containing ~740 bp of the upstream and ~500 bp of the downstream CD1579 homology arms, flanking an *ermB* cassette, was conjugated into 630Δ*erm* using 15 μg/ml thiamphenicol for plasmid selection and 100 μg/ml kanamycin for counterselection of *E. coli*. Faster growing colonies were streaked onto BHIS supplemented with 10 μg/ml thiamphenicol and screened by PCR for upstream or downstream crossover events. Positive colonies were grown in 10 ml BHIS with 100 ng/ml anhydrotetracycline (ATc) and 5 μg/ml erythromycin to induce expression of the CD2517.1 toxin and cure the plasmid. After 24 hours of growth, 2 μl of this culture was streaked on BHIS agar supplemented with 5 μg/ml erythromycin, and colonies were PCR verified for homologous recombination and thiamphenicol sensitivity (**Fig. S1B**).

The *CD1492-H668A* (pMC1000) and *CD2492-H664A* (pMC681) alleles were synthesized and cloned into pMC123 and pUC19, respectively, by Genscript (Piscataway, NJ). The Benchling CRISPR Guide RNA Design tool was used to create a sgRNA targeting *CD1949* (23). The details of vector construction are in the Supplementary Data (**Fig. S2**).

### Sporulation assays and phase contrast microscopy

*C. difficile* strains were grown overnight in BHIS supplemented with 0.1% taurocholate, to promote spore germination, and 0.2% fructose, to inhibit sporulation. In the morning, cells were diluted slightly into BHIS medium and grown until mid-exponential phase (defined as an optical density [OD_600_] of approximately 0.5). Sporulation was examined on 70:30 agar plates or from BHI broth, independently. Aliquots of 0.25 ml were either spread as a lawn onto 70:30 agar supplemented with 2 μg/ml thiamphenicol (19) or diluted 1:10 into BHI broth. Ethanol-resistant sporulation assays were performed after 24 h of growth (H_24_) on 70:30 agar or after three days of growth in BHI broth (H_72_), as previously described (15). Cells were either collected from 70:30 agar and suspended in BHIS medium to an OD_600_ of approximately 1.0 or taken directly from BHI broth. Vegetative cell counts were determined by immediately serially diluting and plating suspended cells onto BHIS. At the same time, ethanol-resistant spore numbers were ascertained by mixing a 0.5 ml aliquot of resuspended cells with 0.3 ml ethanol and 0.2 ml dH_2_O to a final concentration of 28.5% ethanol. This mixture was vortexed and incubated for 15 min to eliminate all vegetative cells. Ethanol-treated cells were serially diluted in 1X PBS containing 0.1% taurocholate and plated onto BHIS with 0.1% taurocholate. Colony forming units (CFU) were enumerated after at least 36 h of growth, and the sporulation frequency was calculated as the total number of spores divided by the total number of spores and vegetative cells. A *spo0A* mutant (MC310) was used as a negative control for sporulation and vegetative cell death. The results represent the means and standard error of the means for at least three independent biological replicates. Statistical significance was performed using a one-way ANOVA, followed by Dunnett’s multiple-comparison test (GraphPad Prism v8.3). Phase contrast microscopy was performed at H_24_ or H_72_, using the resuspended cells, with a Ph3 oil immersion objective on a Nikon Eclipse Ci-L microscope, and at least two fields of view were captured with a DS-Fi2 camera from at least three independent experiments.

### Quantitative reverse transcription PCR analysis (qRT-PCR)

*C. difficile* were cultured on 70:30 agar as a lawn as described above. Cells were collected at H_12_, suspended in 6 ml 1:1:2 ethanol:acetone:water solution, and stored at −80°C. RNA was isolated and subsequently DNase I treated (Ambion) as previously described (24–26). cDNA was synthesized (Bioline) using random hexamers (26). Quantitative real time-PCR (qRT-PCR) analysis was performed in triplicate on 50 ng cDNA using the SensiFAST SYBR & Fluorescein kit (Bioline) and a Roche Lightcycler 96. The results were calculated by the comparative cycle threshold method (27), were normalized to the *rpoC* transcript, and represent the means and standard errors of the means for at least three independent biological replicates. Statistical significance was performed using a one-way ANOVA, followed by a Dunnett’s multiple-comparison test (GraphPad Prism v6.0).

### Enzyme-linked immunosorbent assay (ELISA)

Quantification of TcdA and TcdB toxin present in culture supernatants were performed on *C. difficile* cultures grown in TY medium, pH 7.4 at H_24_, as previously described (28). Briefly, cultures were pelleted, and the supernatants, diluted with the provided dilution buffer, were assayed in technical duplicates using the tgcBIOMICS kit for simultaneous detection of *C. difficile* toxins A and B, according to the manufacturer’s instructions. The averaged results were normalized to the OD_600_ of each respective culture at H_24_, and the results are provided as the means and standard errors of the means for three independent biological replicates. Statistical significance was performed using a one-way ANOVA, followed by a Dunnett’s multiple-comparison test (GraphPad Prism v6.0).

### Phos-tag gel electrophoresis and western blotting analysis

*C. difficile* strains were grown on 70:30 sporulation agar and harvested at H_12_. Lysates were prepared as previously described (15); however, 0.1% Phosphatase Inhibitor Cocktail II (Sigma) was included in the lysis buffer to prevent global protein dephosphorylation. Total protein from lysates was quantitated using the Pierce Micro BCA protein assay kit (Thermo Scientific). Prior to gel electrophoresis, lysates were prepared at 4°C with the exception of an additional 630Δ*erm* aliquot that was briefly heated to 99°C before loading to remove any heat-labile phosphates. Approximately 3 μg of total protein was separated by electrophoresis on a precast 12.5% Super-Sep Phos-tag SDS-PAGE gel (Fujifilm Wako Chemicals Inc, USA) at 90 V for 3.5 h at 4°C. Protein was transferred to 0.2 μm nitrocellulose membrane in transfer buffer containing 10% methanol and 0.04% SDS. Western blot analysis was performed using mouse anti-Spo0A (19) as the primary antibody and goat anti-mouse conjugated with Alexa 488 (Invitrogen) as the secondary antibody. Imaging and densitometry were performed with a ChemiDoc and Image Lab software (Bio-Rad) respectively for three independent experiments.

## RESULTS

### The PtpA (CD1492), PtpB (CD2492) and PtpC (CD1579) orphan histidine kinases inhibit *C. difficile* spore formation

The *C. difficile* 630 genome encodes five orphan histidine kinases, *CD1492*, *CD2492*, *CD1579*, *CD1949*, and *CD1352* which contain conserved catalytic domains that share similarity to *Bacillus* sp. sporulation-associated kinases (16, 29). The CD1352 kinase (CprK) governs a lantibiotic-responsive transporter with no sporulation phenotype and thus, was not included in this study (30). A previous study found that disruption of *CD2492* resulted in decreased sporulation frequency while *in vitro* studies suggested that CD1579 directly phosphorylated Spo0A (16). Our previous work implicated CD1492 as an inhibitor of sporulation, as the sporulation frequency of a *CD1492* mutant was significantly greater than the parent strain (15). To further investigate the impact of these four orphan histidine kinases in *C. difficile* sporulation, we recreated the previously published *CD2492* mutant. We retargeted the group II intron from pCE240 utilizing the same *CD2492* targeting site used by Underwood *et al*. to create a *CD2492* mutant (referred to as the *ptpB* mutant). In addition, we created a *CD1492 CD2492* double mutant (referred to as the *ptpA ptpB* mutant) by introducing the *CD2492*-targeted group II intron into the *CD1492* background (see Materials and Methods for details and the PCR confirmation in **Fig. S1**; the original *CD1492* mutant is referred to as the *ptpA* mutant herein).

We assessed the sporulation phenotypes of the *ptpA*, *ptpB* and *ptpA ptpB* mutants by enumerating ethanol-resistant spores and vegetative cells after 24 h of growth on 70:30 sporulation agar. In these conditions, the 630Δ*erm* parent sporulated at a frequency of ~15.5%. As previously observed, the *ptpA* mutant exhibited a high sporulation frequency of 84.4% (15; Fig. 1A). The *ptpB* mutant and the *ptpA ptpB* double mutant exhibited the same high sporulation frequencies as the *ptpA* mutant (83.6% and 83.7%, respectively; **Fig. 1A**), indicating that the individual genes do not have an additive impact on sporulation. These results suggest that PtpA and PtpB function in the same regulatory pathway to inhibit spore formation. These sporulation phenotypes are also apparent by phase contrast microscopy, as more phase-bright spores were visible in the *ptpA*, *ptpB* and *ptpA ptpB* mutants compared to the parent strain (**Fig. 1B**).

**Figure 1.**
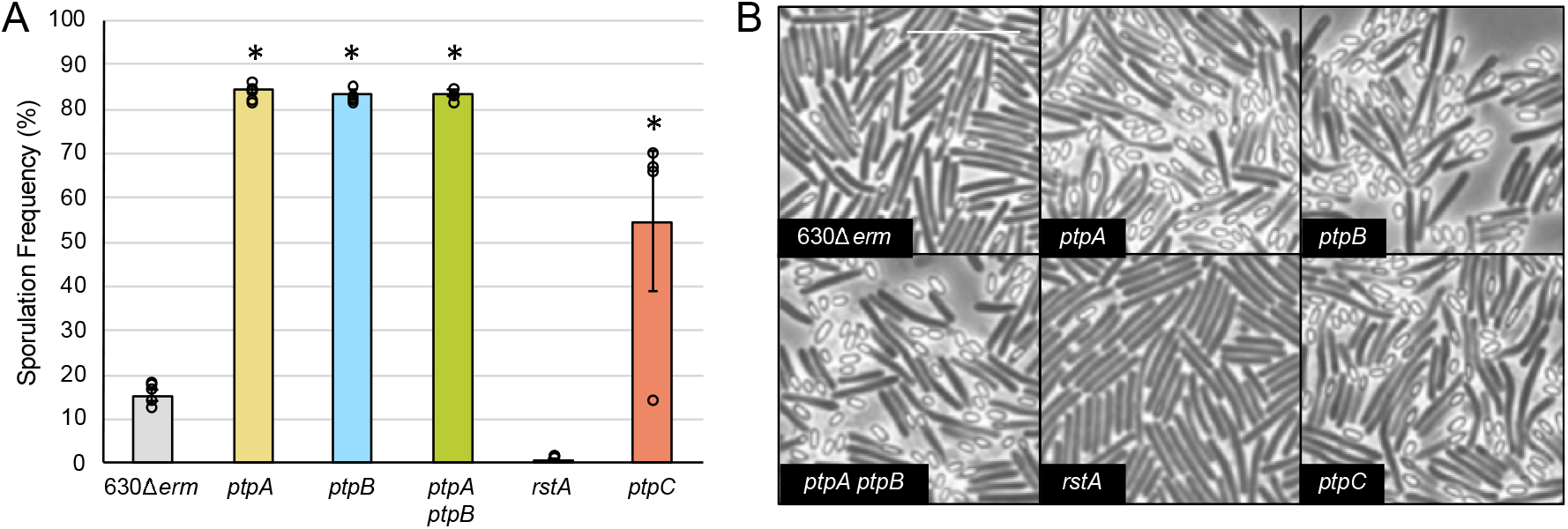
The phosphotransfer proteins PtpA (CD1492), PtpB (CD2492) and PtpC (CD1579) inhibit *C. difficile* spore formation. (**A**) Ethanol-resistant spore formation and (**B**) representative phase contrast micrographs of 630Δ*erm*, *ptpA* (MC674), *ptpB* (MC788), *ptpA ptpB* (MC802), *rstA* (MC1118), and *ptpC* (MC1646) grown on 70:30 sporulation agar at H_24_ (defined as 24 h growth on plates). Sporulation frequency is calculated as the number of ethanol-resistant spores divided by the total number of spores and vegetative cells enumerated. The white scale bar represents 1 μm. *, *P* ≤ 0.01 by a one-way ANOVA followed by Dunnett’s multiple comparisons test.

Based on previous results, we hypothesized that the activity of RstA, a multifunctional regulator that positively influences sporulation (31), may be linked to PtpA (CD1492) activity, as the gene expression profiles and sporulation phenotypes of *rstA* and *ptpA* mutants are opposite (15). Because of both this inverse correlation and the observation that the *ptpB* and *ptpA ptpB* mutants phenocopy the *ptpA* mutant, we included the *rstA* mutant in this study as a comparator. As previously observed, the *rstA* mutant exhibited a significantly low sporulation frequency compared to the parent (**Fig. 1A and B**).

To investigate the impact of PtpC (CD1579) on *C. difficile* sporulation, we created a clean deletion using allele-coupled exchange with a toxin-antitoxin system as a counter-selectable marker to select for plasmid excision (22). Sporulation frequency in the *CD1579* mutant was variable, but ~3.5-fold greater than the 630Δ*erm* parent at 54.8% (**Fig. 1A, B**). This result was somewhat surprising given that PtpC was previously shown to directly phosphorylate Spo0A *in vitro* (16); however, it is not uncommon to observe both kinase and phosphatase activity *in vitro*, even though one direction of phosphate flow is preferred *in vivo* (9). These data suggest that PtpC also inhibits *C. difficile* sporulation, but may not be in the primary regulatory pathway controlling Spo0A dephosphorylation in the conditions tested.

Notably, the sporulation phenotype we observed in the *ptpB* mutant is the opposite of previously published results (16). When Underwood *et al*. created the original *ptpB* mutant, the sporulation frequencies were calculated after 72 h of growth in BHI broth by directly counting carbol fuchsin and malachite green-stained bright-field micrographs. No additional experiments were performed to further probe the sporulation phenotype in the *ptpB* mutant, nor were complementation studies performed (16). We asked whether the *ptpA*, *ptpB,* and *ptpA ptpB* mutants exhibit an alternative sporulation phenotype under different growth conditions. We replicated the sporulation assays performed in BHI medium; however, to quantitate sporulation efficiency, we used the standard ethanol-resistance sporulation assays to enumerate spores, and we assessed sporulation by phase contrast microscopy. The *ptpA*, *ptpB,* and *ptpA ptpB* mutants all hypersporulated in BHI medium (**Fig. 2A**), similar to the sporulation phenotypes observed on 70:30 sporulation agar. Due to the significant amount of cell lysis observed in the phase contract micrographs (**Fig. 2B**), vegetative cells could not be accurately enumerated at this time point from BHI cultures. Thus, the sporulation frequency was counted as spores per ml of culture. These data suggest that the original *ptpB* (low sporulation) phenotype observed by Underwood *et al*. was inaccurate, likely due to the significant cell lysis present after 72 h in BHI. Altogether, our data demonstrate that PtpA and PtpB inhibit *C. difficile* sporulation.

**Figure 2.**
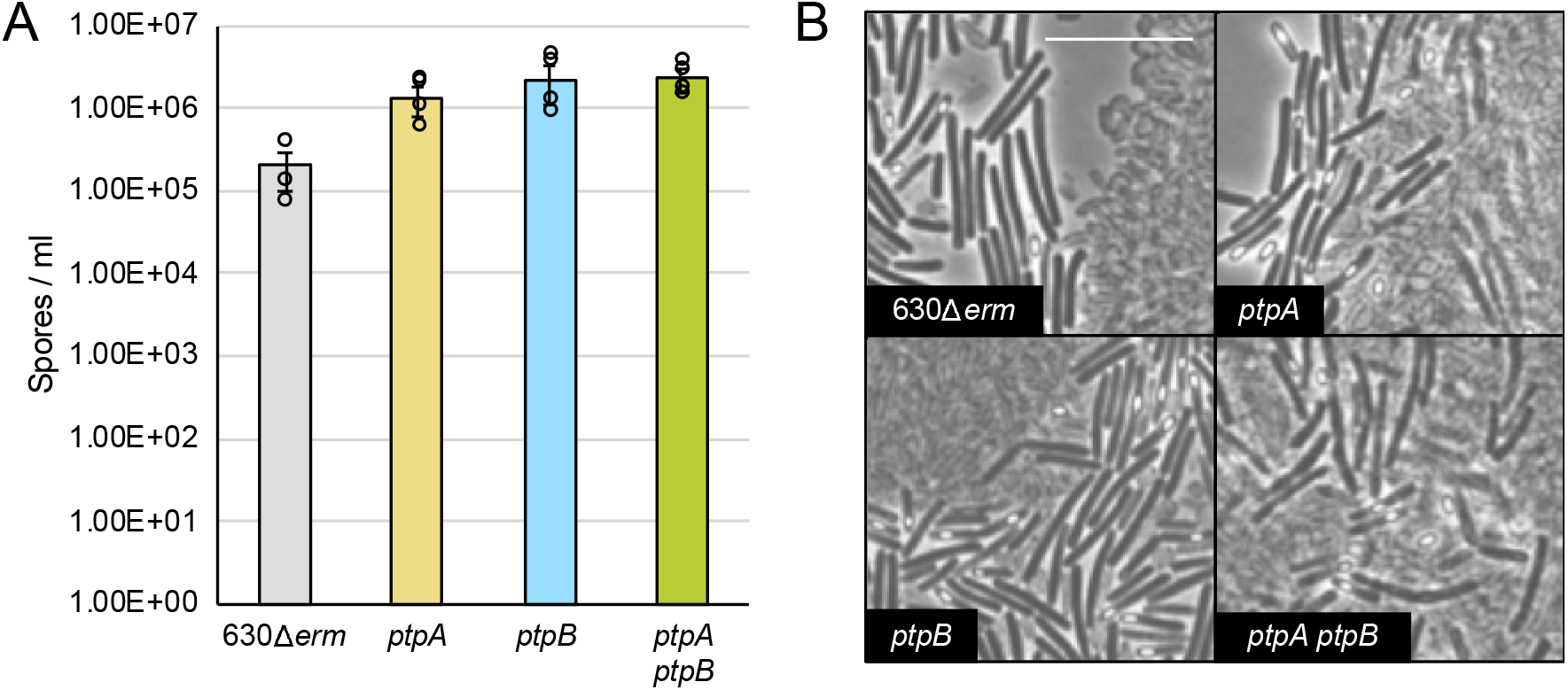
The sporulation frequencies of the *ptpA* (*CD1492)*, *ptpB* (*CD2492)* and *ptpA ptpB* (*CD1492 CD2492)* mutants are increased in BHI medium compared to the parent strain. (**A**) Ethanol-resistant spore formation and (**B**) representative phase contrast micrographs of 630Δ*erm*, *ptpA* (MC674), *ptpB* (MC788) and *ptpA ptpB* (MC802) grown in BHI medium at H_72_. The means and standard errors of the means for three biological replicates are shown. The white scale bar represents 1 μm. No statistical significance observed via a one-way ANOVA followed by Dunnett’s multiple comparisons test.

### CRISPR *i* knockdown of *CD1949* expression does not affect *C. difficile* spore formation

We next asked whether the orphan histidine kinase CD1949 contributes to *C. difficile* sporulation. After numerous unsuccessful attempts to create a *CD1949* null mutant, we utilized the CRISPR interference (CRISPR*i*) tool recently adapted for *C. difficile*, to directly repress *CD1949* transcription (23). Here, the addition of xylose to the medium induced expression of the *dCas9* gene, which encodes a nuclease-deactivated version of caspase-9. dCas9 is then guided to the target transcript by a gene-specific single guide RNA (sgRNA) and subsequently blocks gene transcription. We constructed two different *CD1949*-specific sgRNAs and expressed these in the 630Δ*erm* background. A previously published scrambled sgRNA (sgRNA*-neg)* was included as a control (23). No difference in sporulation frequencies was observed between strains containing the sgRNA-*CD1949* targets compared to the sgRNA-*neg*-containing strain grown on sporulation agar, with or without xylose (**Fig. 3A**). To ensure that *CD1949* was directly targeted by our sgRNA constructs, we measured *CD1949* transcripts using qRT-PCR. *CD1949* transcripts were decreased by ~10-fold or ~20-fold using sgRNA-*C1949-1* or sgRNA-*CD1949-3*, respectively (**Fig. 3B**). These data suggest that CD1949 does not play a role in controlling *C. difficile* sporulation.

**Figure 3.**
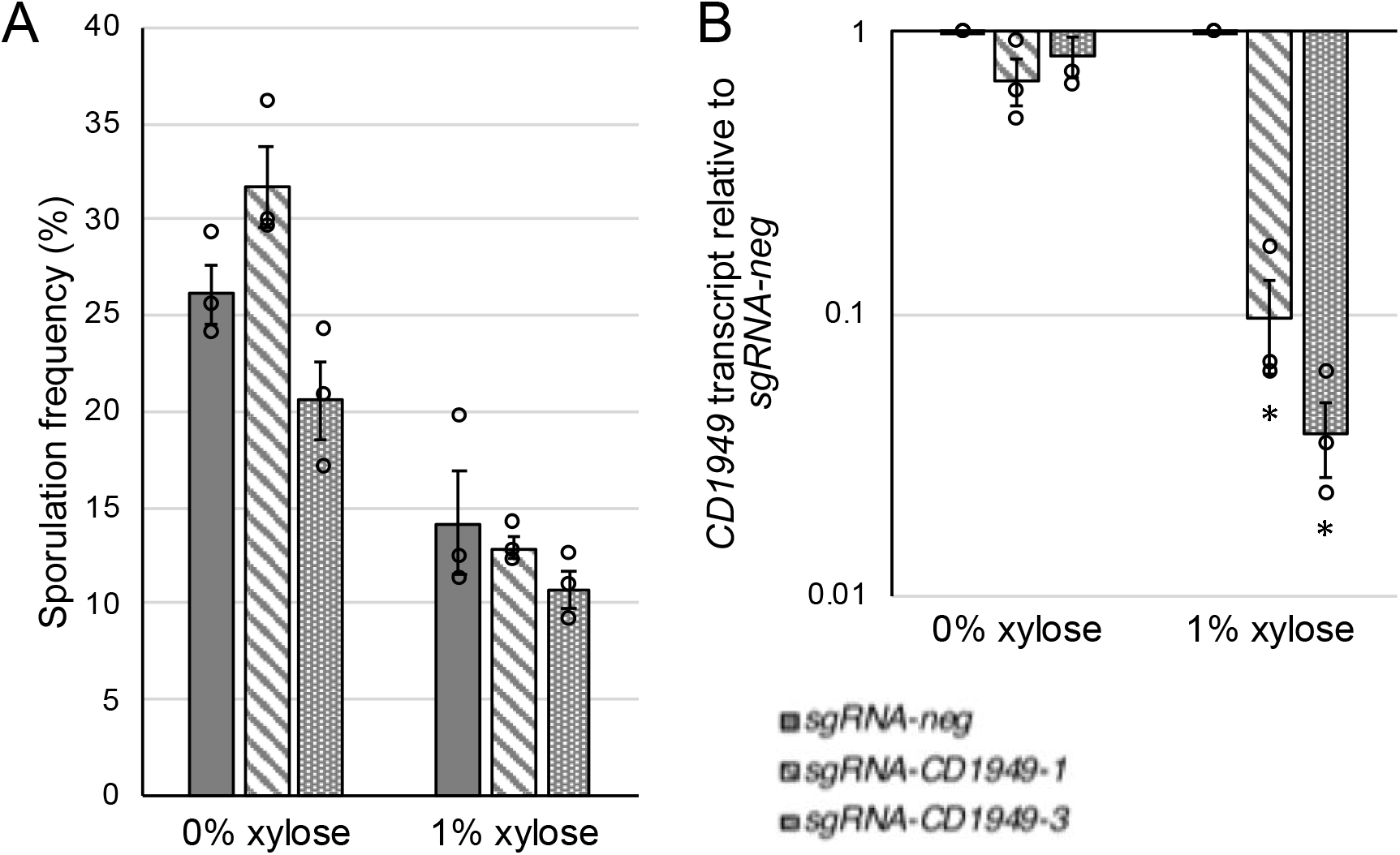
CRISPR *i* knockdown of *CD1949* gene expression does not affect sporulation frequency. (**A**) Ethanol-resistant spore formation at H_24_ and (**B**) qRT-PCR analysis of *CD1949* transcript levels at H_12_ in 630Δ*erm* strains expressing either a scrambled single guide RNA (sgRNA*-neg*) or sgRNAs targeting *CD1949 (*sgRNA*-CD1949-1 and -3*) grown on 70:30 agar supplemented with thiamphenicol 2 μg/ml +/− 1% xylose. Sporulation frequency is calculated as the number of ethanol-resistant spores divided by the total number of spores and vegetative cells enumerated. The means and standard errors of the means for three biological replicates are shown. *, *P* ≤ 0.001 by a one-way ANOVA followed by Dunnett’s multiple comparisons test.

### Deletion of *ptpA* (*CD1492*) and *ptpB* (*CD2492*) results in increased sporulation-specific gene expression and Spo0A activation

To further characterize the sporulation phenotypes of the phosphotransfer protein mutants, we utilized qRT-PCR to measure transcript levels of sporulation-specific genes during the initiation of sporulation at 12 h of growth on sporulation agar (H_12_). We examined expression of *sigF,* encoding the early sporulation forespore-specific sigma factor, *sigE*, which encodes the early mother cell-specific sigma factor, and *spo0A*. The *ptpA*, *ptpB*, and *ptpA ptpB* mutants all presented similarly increased *sigF* (~2.1-2.4-fold) and *sigE* (~1.8-2.1-fold) transcript levels (**Fig. 4A**). As previously observed, the *rstA* mutant had fewer *sigF*, *sigE* and *spo0A* transcripts compared to the parent strain (31; Fig. 4A). Although the *ptpC* mutant had a higher sporulation frequency than the 630Δ*erm* parent at H_24_, sporulation-specific gene expression was not significantly increased by H_12_.

**Figure 4.**
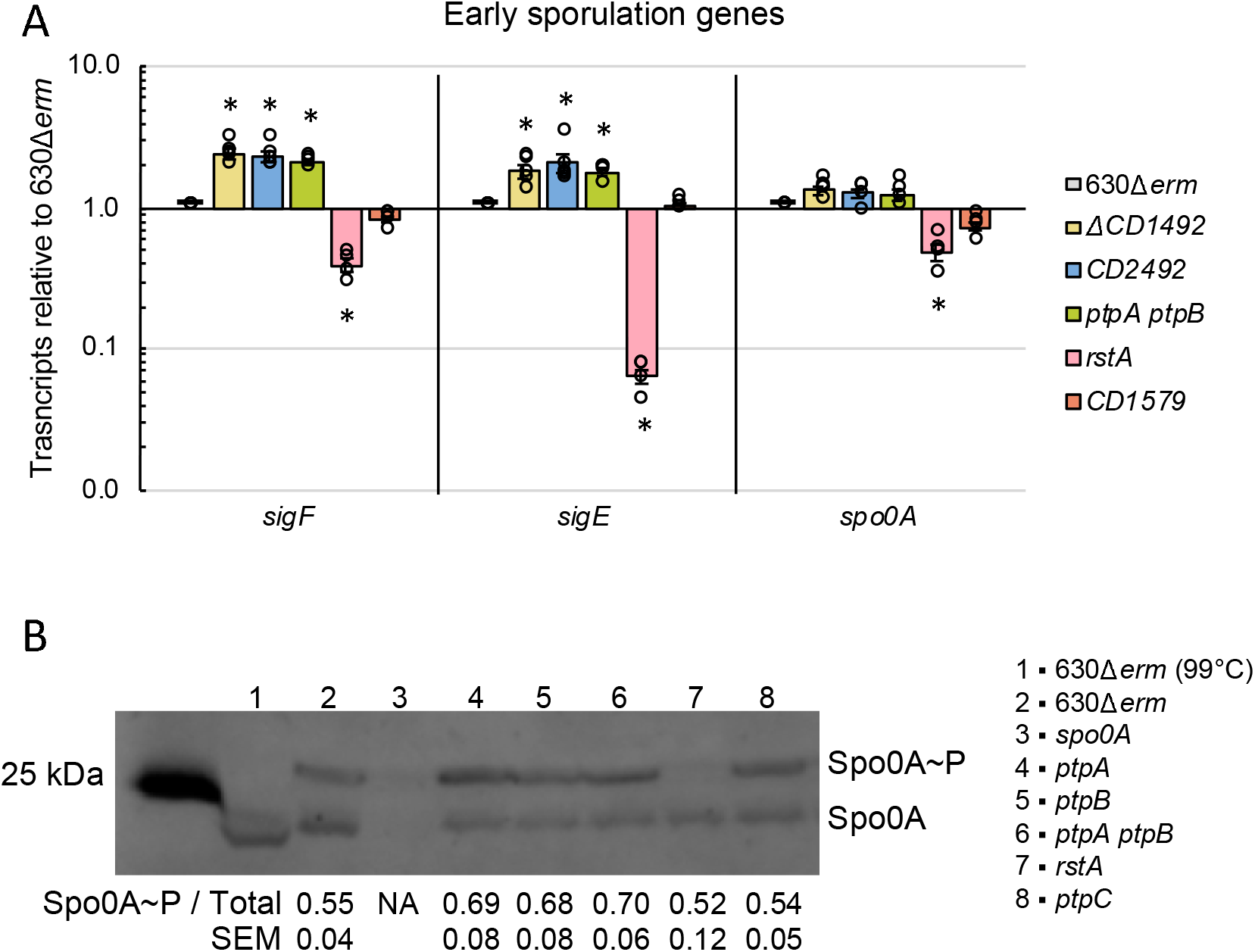
Spo0A-dependent gene expression and Spo0A activation in phosphotransfer protein mutants correlate with endpoint sporulation frequency. (**A**) qRT-PCR analyses of *sigE*, *sigF* and *spo0A* transcripts and (**B**) anti-Spo0A western blot (brightness adjusted) after phos-tag gel separation of unphosphorylated and phosphorylated Spo0A (Spo0A~P) species in 630Δ*erm*, *ptpA* (*CD1492*; MC674), *ptpB* (*CD2492*; MC788), *ptpA ptpB* (*CD1492 CD2492;* MC802), *rstA* (MC1118), and *ptpC* (*CD1579*; MC1646) grown on 70:30 sporulation agar at H_12_. The means and standard errors of the means for at least three biological replicates are shown. *, *P* ≤ 0.05 by a one-way ANOVA followed by Dunnett’s multiple comparisons test.

To determine whether Spo0A phosphorylation was affected during early sporulation in the phosphotransfer protein mutants, we employed phos-tag SDS-polyacrylamide gel electrophoresis. Here, total protein, harvested from cells after 12 h of growth on sporulation agar, was resolved by phos-tag gel electrophoresis. The unphosphorylated (Spo0A) and phosphorylated (Spo0A~P) forms were then detected by western blot with Spo0A antibody. As a control, an aliquot of 630Δ*erm* lysate was boiled to remove any heat-labile phosphate modifications. The protein representing the upper band is the Spo0A~P species, as evidenced by the loss of this upper band after heating (**Fig. 4B**, lane 1 compared to lane 2). There was an increase in the ratio of Spo0A~P to Spo0A in the *ptpA*, *ptpB*, *ptpA ptpB*, and *ptpC* mutants, confirming that a greater proportion of Spo0A protein was phosphorylated in these mutants, corresponding to the onset of sporulation (**Fig. 4B**). Likewise, a much lower ratio of Spo0A~P to Spo0A was observed in the *rstA* mutant, also correlating with the decreased sporulation-specific gene expression and lower sporulation frequency observed in this mutant. Altogether, these data corroborate that PtpA, PtpB, PtpC and RstA all affect Spo0A phosphorylation and thus, early sporulation events in *C. difficile*.

### PtpA (CD1492) and PtpB (CD2492) promote TcdA production

Our previous work demonstrated that the *ptpA* mutant had a ~2-fold decrease in *tcdA* transcript and TcdA protein levels compared to the 630Δ*erm* parent (15). We observed no change in *tcdB* transcript levels in the *ptpA* strain. To determine whether PtpB and PtpC impact toxin production, we measured *tcdA*, *tcdB* and *tcdR* transcript levels in cells grown on 70:30 sporulation agar at H_12_ using qRT-PCR. As we observed previously (15), the *ptpA* mutant exhibited a ~2-fold decrease in *tcdA* transcript levels, but no significant change in *tcdR* or *tcdB* transcripts was observed (**Fig. 5A**). The *ptpB* and *ptpA ptpB* mutants mirrored the changes in toxin transcripts seen in the *ptpA* mutant, exhibiting an ~2-fold decrease in *tcdA* transcript levels, with no effect on *tcdR* and *tcdB* transcript levels (**Fig. 5A**). Toxin transcript levels were not greatly impacted by the loss of *ptpC*, and as we previously observed, the absence of *rstA* resulted in significantly increased *tcdR*, *tcdA* and *tcdB* transcripts, as RstA is a direct repressor of toxin gene transcription (31, 32).

**Figure 5.**
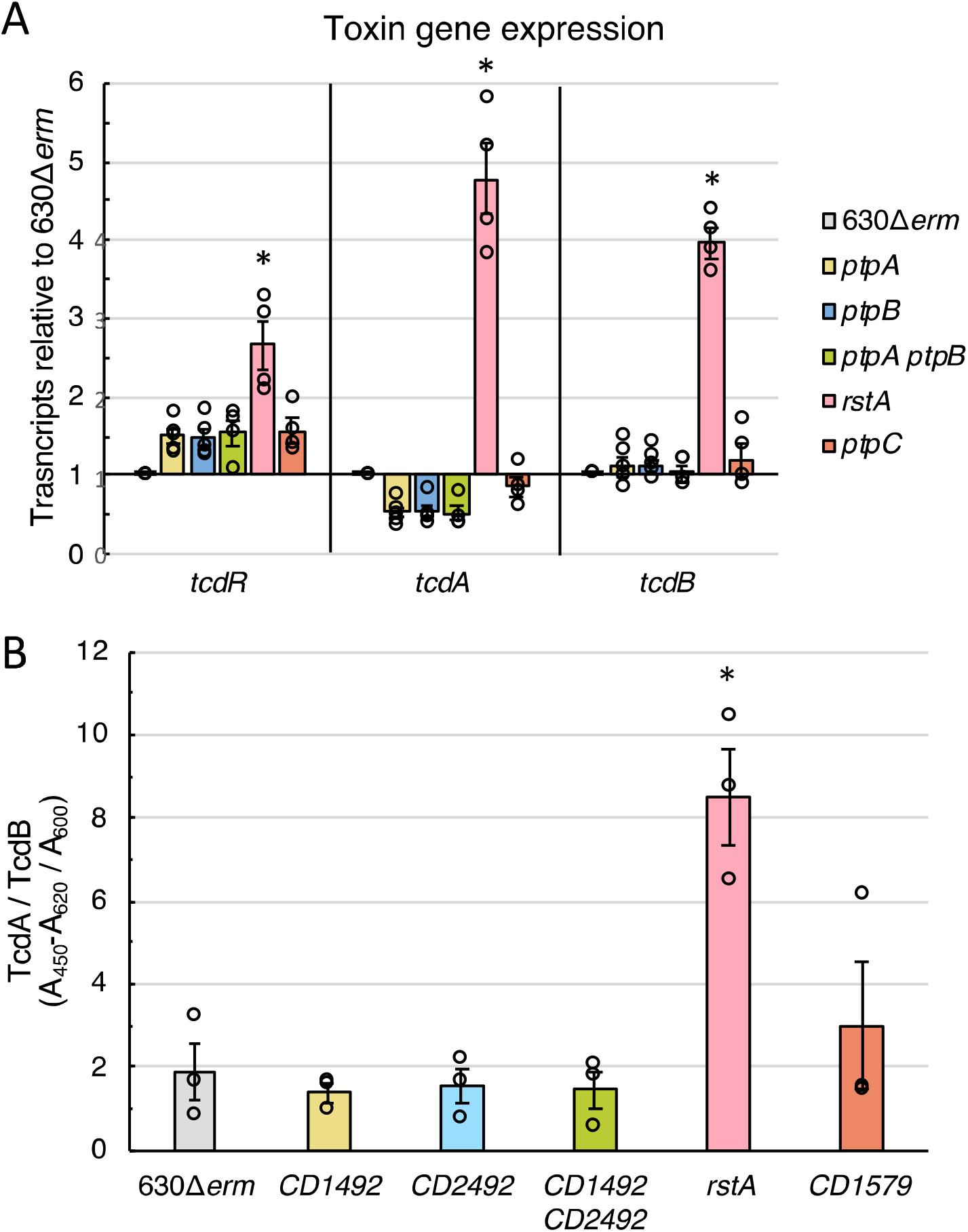
PtpA (CD1492) and PtpB (CD2492) promote TcdA production. (**A**) qRT-PCR analyses of *tcdR*, *tcdA* and *tcdB* transcript levels at H_12_ in 70:30 sporulation agar and (**B**) ELISA analysis of TcdA and TcdB present in the supernatant at H_24_ in TY medium in 630Δ*erm*, *ptpA* (MC674), *ptpB* (MC788), *ptpA ptpB* (MC802), *rstA* (MC1118), and *ptpC* (MC1646). The means and standard errors of the means for at least three biological replicates are shown. *, *P* ≤ 0.05 by a one-way ANOVA followed by Dunnett’s multiple comparisons test.

To further understand the impact that the phosphotransfer proteins exert on toxin production, we measured TcdA and TcdB present in the supernatants of the *ptp* mutants after 24 h growth in TY medium. There was a slight decrease in total toxin production in the *ptpA, ptpB* and *ptpA ptpB* mutants, but this effect was not statistically significant (**Fig. 5B**). Considering the qRT-PCR data, it is likely that the wild-type levels of *tcdB* transcription in the mutants offset the decrease in *tcdA* transcription. Since this ELISA measures the presence of both toxins, the unchanged levels of TcdB in these mutants may mask the repression of TcdA production.

Similar to the variable increase in sporulation frequency in the *ptpC* mutant, we also observed variable concentrations of total TcdA and TcdB toxin present in the supernatant (**Fig. 5B**), suggesting that PtpC does not play a primary role in *C. difficile* toxin production. As expected, TcdA and TcdB toxin were significantly increased in the *rstA* mutant supernatant (31, 32; Fig. 5B).

Although the decrease in *tcdA* transcripts and TcdA/TcdB toxin production in the *ptp* single and double mutants are not statistically significant in this study, we previously found that the decreased TcdA production in the *ptpA* mutant resulted in decreased virulence in the hamster model of infection (15). These data suggest that both PtpA and PtpB enhance *C. difficile* virulence by indirectly promoting *tcdA* transcription through an unknown mechanism. Further, these data comparing the single mutants to the double mutant provide additional support that PtpA and PtpB function in the same regulatory pathway to influence *C. difficile* physiological processes, as the toxin phenotypes in the single and double mutants are all identical.

### PtpA (CD1492) and PtpB (CD2492) are both required for repression of *C. difficile* sporulation, but the conserved histidine residue is not required for PtpB (CD2492) function

To ensure that the sporulation phenotypes exhibited by the *ptpA*, *ptpB*, *ptpA ptpB,* and *ptpC* mutants were due to disruption or loss of the targeted gene, we complemented these mutants by expressing each locus under the control of its native promoter on an exogenous plasmid. Expression of *ptpA* or *ptpB* from their native promoters restored the *ptpA* and *ptpB* single mutants’ sporulation frequencies to below wild-type levels (**Fig. 6A**). However, expressing *ptpA* in the *ptpB* mutant or *ptpB* in the *ptpA* mutant did not complement sporulation, further supporting that PtpA and PtpB functions are not redundant (**Fig. 6A**). Complementation of the *CD1492 CD2492* double mutant required the expression of both *ptpA* and *ptpB*; expression of a single phosphotransfer protein in the double mutant was not enough to exert any impact on sporulation frequency (**Fig. 6A**). Altogether, these data indicate that PtpA and PtpB function together in a regulatory pathway to inhibit spore formation and that their functions are not interchangeable.

**Figure 6.**
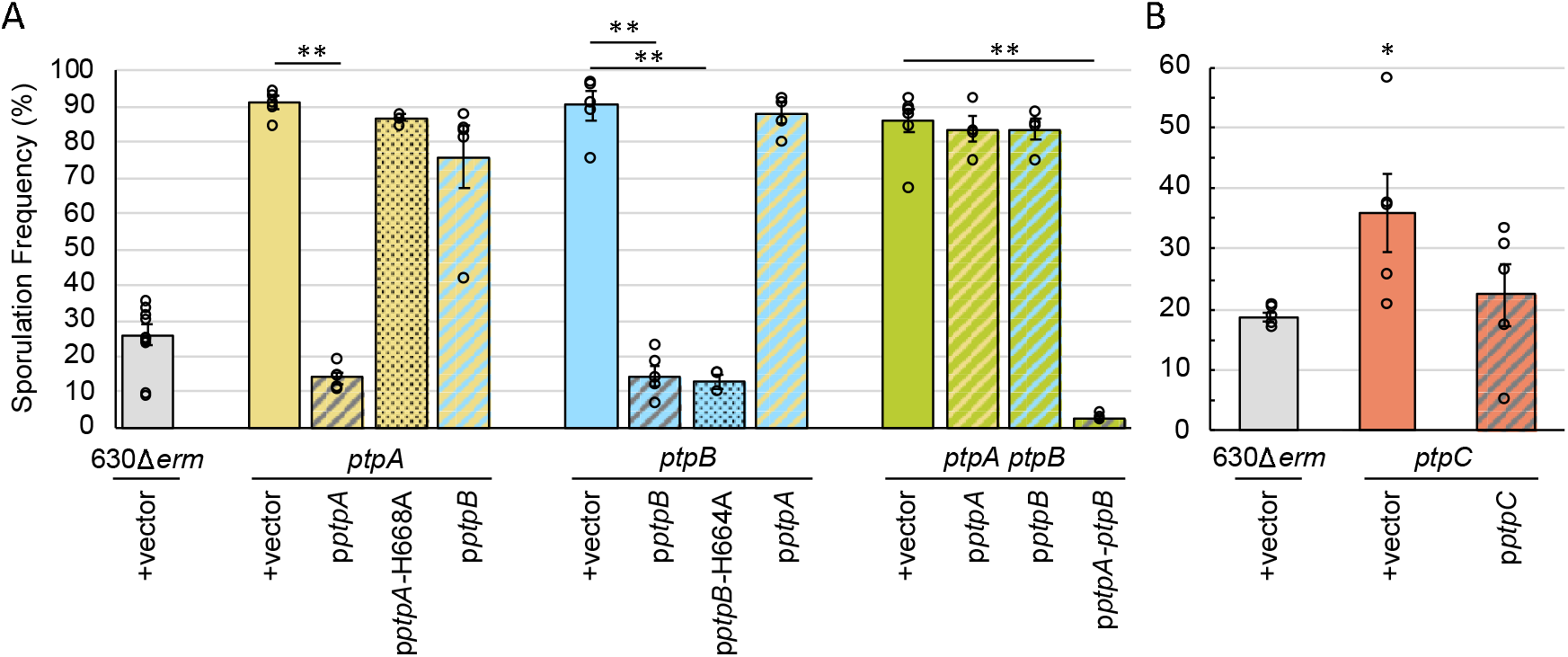
PtpA (CD1492) and PtpB (CD2492) are both required for repression of *C. difficile* sporulation, but the conserved histidine residue is not required for PtpB (CD2492) function. Ethanol-resistant spore formation in (**A**) 630Δ*erm* pMC123 (MC324) *ptpA* pMC123 (MC964), *ptpA* p*ptpA* (MC998), *ptpA* p*ptpA*-H668A (MC1812), *ptpA* p*ptpB* (MC965), *ptpB* pMC123 (MC966), *ptpB* p*ptpB* (MC967), *ptpB* p*ptpB-H664A* (MC1030), *ptpB* p*ptpA* (MC999), *ptpA ptpB* pMC123 (MC968), *ptpA ptpB* p*ptpA* (MC1000), *ptpA ptpB* p*ptpB* (MC969), and *ptpA ptpB* p*ptpA-ptpB* (MC1396) and (**B**) 630Δ*erm* pMC123 (MC324), *ptpC* pMC123 (MC1672), and *ptpC* p*ptpC* (MC1673) grown on 70:30 sporulation agar supplemented with 2 μg/ml thiamphenicol at H_24_. Sporulation frequency is calculated as the number of ethanol-resistant spores divided by the total number of spores and vegetative cells enumerated. The means and standard errors of the means for at least four biological replicates are shown. Note the difference in scales between panels A and B. *, *P* ≤ 0.05; **, *P* ≤ 0.001 by a one-way ANOVA followed by Dunnett’s multiple comparisons test.

The autophosphorylation and phosphotransferase activities of sensor histidine kinases rely on a conserved histidine residue located in the dimerization and histidine phosphotransfer domain (DHpt) (33, 34). These conserved histidine residues are present in PtpA, PtpB, and PtpC and were proposed to be critical for phosphotransfer to an aspartyl residue in Spo0A (16). Replacing the conserved histidine residue with alanine in a histidine kinase disables the autophosphorylation and phosphotransfer activity of the protein, resulting in a nonfunctional protein (35). Our previous work showed that overexpression of *ptpA-H668A* in the *ptpA* background did not reduce sporulation (15), suggesting that the histidine residue is critical for CD1492 function in sporulation. We were able to replicate these results by expressing *ptpA-H668A* from its native promoter, rather than the inducible promoter used previously (**Fig. 6A**). Surprisingly, the corresponding *ptpB-H664A* allele did complement the *ptpB* mutant (**Fig. 6A**), indicating that this histidine residue is not necessary for PtpB to repress sporulation.

To confirm that the *ptpC* mutation was responsible for the hypersporulation phenotype observed, the *ptpC* gene was expressed from its native promoter on an exogenous plasmid. Although the sporulation phenotypes were variable in the *ptpC* strains harboring the control and the p*ptpC* plasmids, we did observe an ~1.6-fold reduction in sporulation frequency in the *ptpC* p*ptpC* complementation strain compared to *ptpC* containing the vector control (**Fig. 6B**). Because of the variable phenotype, we performed whole genome sequencing on the *ptpC* mutant and found no additional mutations besides the replacement of the *ptpC* allele with the *ermB* cassette (data not shown).

### Expression of the *ptp* site-directed mutants result in a dominant negative phenotype

To further probe the function of the conserved histidine residues, we expressed the *ptpA*, *ptpB* and *ptpC* wild-type alleles and histidine site-directed mutations from their native promoters in the 630Δ*erm* background. Comparable to our previous study (15), sporulation frequency decreased when *ptpA* was expressed from its native promoter, compared to the parent strain containing the empty vector (**Fig. 7A, B**; from 33.3% in 630Δ*erm* pMC123 to 14.4% in 630Δ*erm* p*ptpA*). We observed a similar effect when *ptpB* was expressed in 630Δ*erm* (**Fig. 7A, B**; to 16.0% in 630Δ*erm* p*ptpB*), indicating that PtpA and PtpB are able to reduce sporulation in an otherwise wild-type background.

**Figure 7.**
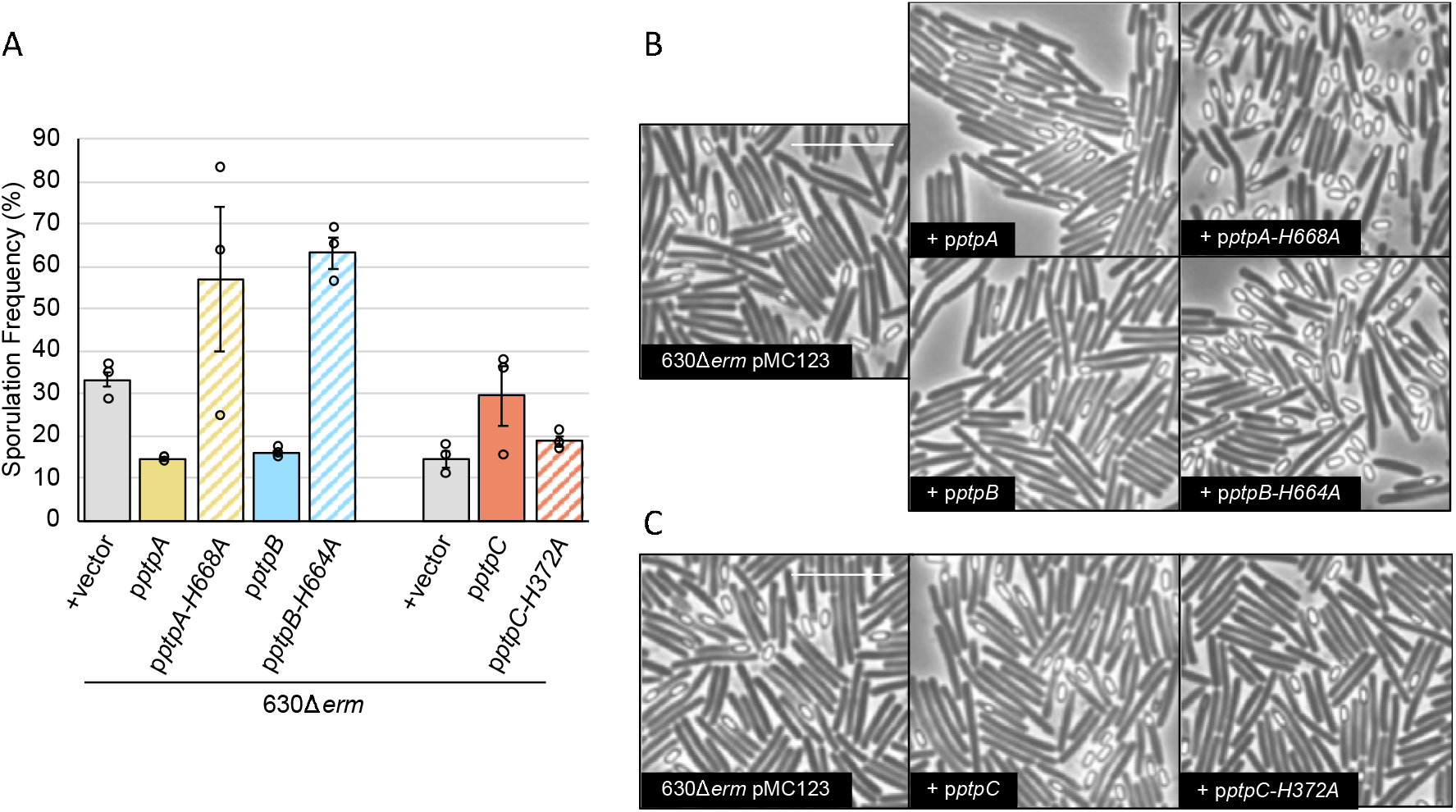
*ptpA* (*CD1492*) and *ptpB* (*CD2492*) expression in the 630Δ*erm* background decreases sporulation frequency, while *ptpA-H668A* and *ptpB-H664A* expression results in a dominant negative phenotype. (**A**) Ethanol-resistant spore formation and representative phase contrast micrographs in (**B, C**) 630Δ*erm* pMC123 (MC324), 630Δ*erm* p*ptpA* (MC2024), 630Δ*erm* p*ptpA*-*H668A* (MC2025), 630Δ*erm* p*ptpB* (MC2026), 630Δ*erm* p*ptpB-H664A* (MC2027), 630Δ*erm* p*ptpC* (MC2030), 630Δ*erm* p*ptpC-H372A* (MC2031) grown on 70:30 sporulation agar supplemented with 2 μg/ml thiamphenicol at H_24_. Experiments with *ptpA* and *ptpB* were performed at different times than with *ptpC*, and the 630Δ*erm* pMC123 (MC324) control strain is shown for each. Sporulation frequency is calculated as the number of ethanol-resistant spores divided by the total number of spores and vegetative cells enumerated. The means and standard errors of the means for three biological replicates are shown. The white scale bar represents 1 μm. *, *P* ≤ 0.05 by a one-way ANOVA followed by Dunnett’s multiple comparisons test comparing each strain to 630Δ*erm* pMC123 (MC324).

Since histidine kinases function as oligomers, we next hypothesized that expression of the nonfunctional *ptpA-H668A* allele in the parental background would result in nonfunctional hetero-oligomers. These hetero-oligomers would be unable to function as a phosphatase, resulting in increased sporulation and a dominant negative phenotype. As predicted, sporulation frequency increased in 630Δ*erm* p*ptpA-H668A* by ~1.7-fold compared to the parent strain (**Fig. 7A, B**). Although the *ptpB-H664A* allele complemented the *ptpB* mutant, we also observed a dominant negative phenotype when p*ptpB-H664A* was expressed in 630Δ*erm*, as sporulation frequency increased ~1.9-fold (**Fig. 7A, B**). In contrast to the complementation study (**Fig. 6A**), these data suggest that the conserved histidine residue of PtpB plays a role in *C. difficile* sporulation.

Finally, we examined the effect on sporulation when *ptpC* is expressed in the 630*erm* background. Although not statistically significant because of variability, spore formation was increased by ~2-fold when the wild-type *ptpC* allele was expressed in 630Δ*erm* (**Fig. 7A, C**). However, sporulation frequency was increased by only ~1.3-fold when the *ptpC-H372A* allele was expressed, suggesting that the histidine residue impacts the ability for PtpC to influence *C. difficile* spore formation.

## DISCUSSION

Although the morphological changes that produce a dormant spore are conserved between Clostridia and the well-studied Bacilli, the regulatory pathway and factors that control sporulation initiation in *C. difficile* are not well defined (10–12). In all spore-forming bacteria, the onset of sporulation is governed by the essential regulator of sporulation, Spo0A, which is activated by phosphorylation and inactivated by dephosphorylation (10, 36). The broadly studied *Bacillus* sp. use an expanded two-component system, known as a phosphorelay, to transfer a phosphate signal from sporulation-associated sensor histidine kinases via two phosphotransfer proteins to Spo0A, to trigger the onset of sporulation. An orthologous expanded sporulation regulatory pathway leading to Spo0A activation is not encoded in the *C. difficile* genome or other Clostridia (7, 10, 12); however, several orphan sensor histidine kinases, including CD1492, CD2492 and CD1579, have been implicated in controlling sporulation initiation by influencing Spo0A phosphorylation (15, 16).

This investigation has expanded what was previously known about how CD1492 (PtpA), CD2492 (PtpB) and CD1579 (PtpC) impact *C. difficile* spore formation (15, 16). The data demonstrate that PtpA and PtpB function in the same regulatory pathway to control Spo0A activation, as the sporulation phenotypes and gene expression profiles of the single and double mutants are identical. However, neither protein can fulfill the role of the other, indicating that PtpA and PtpB functions are nonredundant. Further, the data suggest that these proteins function together, not necessarily stepwise, as neither protein is epistatic to the other. It is possible that PtpA and PtpB form hetero-oligomers and/or do not function as phosphatases without both proteins present. We hypothesize that PtpA and PtpB directly bind to and dephosphorylate Spo0A, although alternatively, PtpA and PtpB may interact with an intermediate factor(s) that directly phosphorylates Spo0A or serve as an endpoint in a serial dephosphorylation pathway. We attempted to assess potential direct protein-protein interactions by using tagged, full-length proteins in previously validated bacterial adenylate cyclase two hybrid (BACTH) and split luciferase assays (37, 38). However, these approaches have been unsuccessful thus far, likely because PtpA and PtpB are membrane proteins that are toxic when expressed in *E. coli,* resulting in unstable constructs. We also unsuccessfully attempted to capture these interactions *in vivo* with co-immunoprecipitation assays using dually-tagged full-length recombinant proteins, followed by western blotting. In future studies, we will employ alternative approaches, such as working with cytosolic PtpA and PtpB truncations (14, 34, 39) and performing co-immunoprecipitation studies followed by mass spectrometry analyses. Identifying the direct binding partners of these phosphotransfer proteins is a high priority.

The role of PtpC in sporulation is less clear. A *ptpC* null mutant had increased, but variable, sporulation. In contradiction, overexpression of *ptpC* in 630Δ*erm* also increased sporulation frequency. But, expression of the *ptpC-H372A* allele had no impact on sporulation. PtpC was previously shown to phosphorylate Spo0A *in vitro*, suggesting that PtpC positively controls *C. difficile* sporulation initiation (16). However, the contribution of the conserved PtpC histidine residue for *in vitro* phosphate transfer to Spo0A was not tested. These data suggest that PtpC has the ability to perform both kinase and phosphatase functions.

The environmental and/or intracellular signals that control the activities of PtpA, PtpB and PtpC are unknown. These proteins may have kinase or phosphatase activity under differing conditions, similar to the well-studied EnvZ histidine kinase of *Escherichia coli* and *Salmonella enterica* (33, 34), the pH-sensing HK853 from *Thermotoga maritima* (40), and the quorum sensing sensor kinases of *Vibrio* sp., LuxN and LuxQ (41, 42). LuxN requires a conserved aspartate residue, but not the conserved histidine residue, for phosphatase activity (42), and EnvZ retains phosphatase activity when several other residues are substituted for the histidine residue (43). Further, another *C. difficile* histidine kinase, CprK, has also exhibited potential phosphatase activity in the absence of its conserved histidine residue (30). Retention of phosphatase activity may explain why the PtpB-H664A mutant remained functional in complementation studies yet displayed a dominant negative phenotype in the parent strain. We hypothesize that the histidine residue is not required for PtpB phosphatase activity, but is required for PtpA activity. Further, PtpA, PtpB, and PtpC all contain a conserved E/DxxT/N motif in which the E/D residue is critical for kinase activity and the T/N motif is necessary for phosphatase activity (44, 45). Along with our data, the presence of this conserved motif provides additional evidence that PtpA, PtpB, and PtpC may possess dual kinase and phosphatase activities. Elucidating the molecular mechanisms by which PtpA, PtpB, and PtpC control phosphate flux to Spo0A are a focus of our future studies.

Regulation of *ptpA*, *ptpB,* and *ptpC* gene expression may also influence the timing of accumulation and activity of these proteins. The transition phase sigma factor SigH directly activates *ptpB* transcription (46), while the inactivation of *sigB*, a general stress response sigma factor, results in decreased *ptpA* expression and increased *ptpC* expression (47). Additionally, the catabolite control protein, CcpA, appears to indirectly repress *ptpC* expression in response to glucose (48). Altogether, the expansive list of global regulators that influence *ptpA*, *ptpB,* and *ptpC* gene expression underscore that the pathways that control Spo0A phosphorylation and dephosphorylation are under complex regulatory control. Understanding when these phosphotransfer proteins are expressed and when they are active will provide insight into what signals *C. difficile* couples to the onset of spore formation.

Our data demonstrate that *C. difficile* utilizes at least two nonredundant pathways to regulate Spo0A activation. This appears in contrast to *B. subtilis*, which positively controls phosphate flux from the kinases to Spo0A through the intermediate proteins, Spo0F and Spo0B. To modulate the phosphate flow to Spo0A, *B. subtilis* employs several aspartyl phosphatases, which directly dephosphorylate Spo0F or Spo0A (49–51), and kinase inhibitors, which prevent KinA autophosphorylation and/or phosphotransfer (52, 53). *C. difficile* also encodes orthologous aspartyl-phosphatases and kinase inhibitor genes (11), potentially providing additional regulatory mechanisms to inhibit sporulation under specific conditions, even if the precise function or target is not conserved with *B. subtilis*. Although the Clostridia are hypothesized to have a simplified Spo0A activation pathway, *C. difficile* and its relatives *Clostridium acetobutylicum*, *Acetivibrio thermocellus,* and *C. perfringens* employ multiple, potentially dual function, histidine kinases to control Spo0A phosphorylation (13, 14, 54, 55). These results suggest that sporulation initiation is more tightly regulated in response to environmental and intracellular cues in Clostridia than previously credited.

As PtpA, PtpB and PtpC all inhibit sporulation, one of the biggest questions remaining is what factors are primarily responsible for Spo0A activation? The multifunctional regulator RstA positively influences Spo0A phosphorylation through an unknown molecular mechanism (28, 31). Although there is no evidence that RstA directly binds Spo0A or functions as a kinase, it remains possible that RstA phosphorylates Spo0A or an intermediate or blocks Spo0A dephosphorylation by steric hindrance. Spo0A phosphorylation may also be controlled directly by unidentified kinases. These potentially unknown kinases are difficult to predict based on knowledge from well-studied systems, as there is low conservation between clostridial or *Bacillus* spore formers. Identifying proteins that directly interact with known sporulation factors may uncover additional regulators that impact sporulation initiation, helping to unravel the regulatory pathways and molecular mechanisms that influence the ability for *C. difficile* transmission and survival.

## ACKNOWLEDGEMENTS

We are grateful to the members of the McBride lab and Joseph Sorg for their helpful suggestions and discussions throughout the course of this work. We are also thankful to Johann Peltier for the gift of pMSR. This research was supported by the U.S. National Institutes of Health through research grants AI116933 and AI156052 to S.M.M and GM008490 to M.A.D. The content of this manuscript is solely the responsibility of the authors and does not necessarily reflect the official views of the National Institutes of Health.

## REFERENCES

1. Ferrari FA, Trach K, LeCoq D, Spence J, Ferrari E, Hoch JA. 1985. Characterization of the spo0A locus and its deduced product. Proc Natl Acad Sci U S A 82:2647-51.

2. Deakin LJ, Clare S, Fagan RP, Dawson LF, Pickard DJ, West MR, Wren BW, Fairweather NF, Dougan G, Lawley TD. 2012. The Clostridium difficile spo0A gene is a persistence and transmission factor. Infect Immun 80:2704-11.

3. Rosenbusch KE, Bakker D, Kuijper EJ, Smits WK. 2012. C. difficile 630Deltaerm Spo0A regulates sporulation, but does not contribute to toxin production, by direct high-affinity binding to target DNA. PLoS One 7:e48608.

4. Hoch JA. 1993. Regulation of the phosphorelay and the initiation of sporulation in Bacillus subtilis. Annu Rev Microbiol 47:441-65.

5. Bird TH, Grimsley JK, Hoch JA, Spiegelman GB. 1993. Phosphorylation of Spo0A activates its stimulation of in vitro transcription from the Bacillus subtilis spoIIG operon. Mol Microbiol 9:741-9.

6. Baldus JM, Green BD, Youngman P, Moran CP, Jr. 1994. Phosphorylation of Bacillus subtilis transcription factor Spo0A stimulates transcription from the spoIIG promoter by enhancing binding to weak 0A boxes. J Bacteriol 176:296-306.

7. Fimlaid KA, Bond JP, Schutz KC, Putnam EE, Leung JM, Lawley TD, Shen A. 2013. Global Analysis of the Sporulation Pathway of Clostridium difficile. PLoS Genet 9:e1003660.

8. Sonenshein AL. 2000. Control of sporulation initiation in Bacillus subtilis. Curr Opin Microbiol 3:561-6.

9. Burbulys D, Trach KA, Hoch JA. 1991. Initiation of sporulation in B. subtilis is controlled by a multicomponent phosphorelay. Cell 64:545-52.

10. Paredes CJ, Alsaker KV, Papoutsakis ET. 2005. A comparative genomic view of clostridial sporulation and physiology. Nat Rev Microbiol 3:969-78.

11. Edwards AN, McBride SM. 2014. Initiation of sporulation in Clostridium difficile: a twist on the classic model. FEMS Microbiol Lett 358:110-8.

12. Shen A, Edwards AN, Sarker MR, Paredes-Sabja D. 2019. Sporulation and Germination in Clostridial Pathogens. Microbiol Spectr 7.

13. Steiner E, Dago AE, Young DI, Heap JT, Minton NP, Hoch JA, Young M. 2011. Multiple orphan histidine kinases interact directly with Spo0A to control the initiation of endospore formation in Clostridium acetobutylicum. Mol Microbiol 80:641-54.

14. Freedman JC, Li J, Mi E, McClane BA. 2019. Identification of an Important Orphan Histidine Kinase for the Initiation of Sporulation and Enterotoxin Production by Clostridium perfringens Type F Strain SM101. mBio 10.

15. Childress KO, Edwards AN, Nawrocki KL, Woods EC, Anderson SE, McBride SM. 2016. The Phosphotransfer Protein CD1492 Represses Sporulation Initiation in Clostridium difficile. Infect Immun doi:10.1128/IAI.00735-16.

16. Underwood S, Guan S, Vijayasubhash V, Baines SD, Graham L, Lewis RJ, Wilcox MH, Stephenson K. 2009. Characterization of the sporulation initiation pathway of Clostridium difficile and its role in toxin production. J Bacteriol 191:7296-305.

17. Edwards AN, Suarez JM, McBride SM. 2013. Culturing and maintaining Clostridium difficile in an anaerobic environment. J Vis Exp doi:10.3791/50787:e50787.

18. Sorg JA, Dineen SS. 2009. Laboratory maintenance of Clostridium difficile. Curr Protoc Microbiol Chapter 9:Unit9A 1.

19. Putnam EE, Nock AM, Lawley TD, Shen A. 2013. SpoIVA and SipL are Clostridium difficile spore morphogenetic proteins. J Bacteriol 195:1214-25.

20. Purcell EB, McKee RW, McBride SM, Waters CM, Tamayo R. 2012. Cyclic diguanylate inversely regulates motility and aggregation in Clostridium difficile. J Bacteriol 194:3307-16.

21. Edwards AN, Williams CL, Pareek N, McBride SM, Tamayo R. 2021. c-di-GMP inhibits early sporulation in Clostridioides difficile. BioRxiv doi:10.1101/2021.06.24.449855.

22. Peltier J, Hamiot A, Garneau JR, Boudry P, Maikova A, Hajnsdorf E, Fortier LC, Dupuy B, Soutourina O. 2020. Type I toxin-antitoxin systems contribute to the maintenance of mobile genetic elements in Clostridioides difficile. Commun Biol 3:718.

23. Muh U, Pannullo AG, Weiss DS, Ellermeier CD. 2019. A Xylose-Inducible Expression System and a CRISPR Interference Plasmid for Targeted Knockdown of Gene Expression in Clostridioides difficile. J Bacteriol 201.

24. McBride SM, Sonenshein AL. 2011. Identification of a genetic locus responsible for antimicrobial peptide resistance in Clostridium difficile. Infect Immun 79:167-76.

25. Dineen SS, McBride SM, Sonenshein AL. 2010. Integration of metabolism and virulence by Clostridium difficile CodY. J Bacteriol 192:5350-62.

26. Edwards AN, Nawrocki KL, McBride SM. 2014. Conserved oligopeptide permeases modulate sporulation initiation in Clostridium difficile. Infect Immun 82:4276-91.

27. Schmittgen TD, Livak KJ. 2008. Analyzing real-time PCR data by the comparative C(T) method. Nat Protoc 3:1101-8.

28. Edwards AN, Krall EG, McBride SM. 2020. Strain-Dependent RstA Regulation of Clostridioides difficile Toxin Production and Sporulation. J Bacteriol 202.

29. Sebaihia M, Wren BW, Mullany P, Fairweather NF, Minton N, Stabler R, Thomson NR, Roberts AP, Cerdeno-Tarraga AM, Wang H, Holden MT, Wright A, Churcher C, Quail MA, Baker S, Bason N, Brooks K, Chillingworth T, Cronin A, Davis P, Dowd L, Fraser A, Feltwell T, Hance Z, Holroyd S, Jagels K, Moule S, Mungall K, Price C, Rabbinowitsch E, Sharp S, Simmonds M, Stevens K, Unwin L, Whithead S, Dupuy B, Dougan G, Barrell B, Parkhill J. 2006. The multidrug-resistant human pathogen Clostridium difficile has a highly mobile, mosaic genome. Nat Genet 38:779-86.

30. Suarez JM, Edwards AN, McBride SM. 2013. The Clostridium difficile cpr locus is regulated by a noncontiguous two-component system in response to type A and B lantibiotics. J Bacteriol 195:2621-31.

31. Edwards AN, Tamayo R, McBride SM. 2016. A novel regulator controls Clostridium difficile sporulation, motility and toxin production. Mol Microbiol 100:954-71.

32. Edwards AN, Anjuwon-Foster BR, McBride SM. 2019. RstA Is a Major Regulator of Clostridioides difficile Toxin Production and Motility. MBio 10.

33. Igo MM, Ninfa AJ, Stock JB, Silhavy TJ. 1989. Phosphorylation and dephosphorylation of a bacterial transcriptional activator by a transmembrane receptor. Genes Dev 3:1725-34.

34. Dutta R, Inouye M. 1996. Reverse phosphotransfer from OmpR to EnvZ in a kinase-/phosphatase+ mutant of EnvZ (EnvZ.N347D), a bifunctional signal transducer of Escherichia coli. J Biol Chem 271:1424-9.

35. Hoch JA. 2000. Two-component and phosphorelay signal transduction. Curr Opin Microbiol 3:165-70.

36. Brown DP, Ganova-Raeva L, Green BD, Wilkinson SR, Young M, Youngman P. 1994. Characterization of spo0A homologues in diverse Bacillus and Clostridium species identifies a probable DNA-binding domain. Mol Microbiol 14:411-26.

37. Karimova G, Pidoux J, Ullmann A, Ladant D. 1998. A bacterial two-hybrid system based on a reconstituted signal transduction pathway. Proc Natl Acad Sci U S A 95:5752-6.

38. Oliveira Paiva AM, Friggen AH, Qin L, Douwes R, Dame RT, Smits WK. 2019. The Bacterial Chromatin Protein HupA Can Remodel DNA and Associates with the Nucleoid in Clostridium difficile. J Mol Biol 431:653-672.

39. Goodman AL, Merighi M, Hyodo M, Ventre I, Filloux A, Lory S. 2009. Direct interaction between sensor kinase proteins mediates acute and chronic disease phenotypes in a bacterial pathogen. Genes Dev 23:249-59.

40. Liu Y, Rose J, Huang S, Hu Y, Wu Q, Wang D, Li C, Liu M, Zhou P, Jiang L. 2017. A pH-gated conformational switch regulates the phosphatase activity of bifunctional HisKA-family histidine kinases. Nat Commun 8:2104.

41. Freeman JA, Bassler BL. 1999. A genetic analysis of the function of LuxO, a two-component response regulator involved in quorum sensing in Vibrio harveyi. Mol Microbiol 31:665-77.

42. Freeman JA, Lilley BN, Bassler BL. 2000. A genetic analysis of the functions of LuxN: a two-component hybrid sensor kinase that regulates quorum sensing in Vibrio harveyi. Mol Microbiol 35:139-49.

43. Hsing W, Silhavy TJ. 1997. Function of conserved histidine-243 in phosphatase activity of EnvZ, the sensor for porin osmoregulation in Escherichia coli. J Bacteriol 179:3729-35.

44. Huynh TN, Noriega CE, Stewart V. 2010. Conserved mechanism for sensor phosphatase control of two-component signaling revealed in the nitrate sensor NarX. Proc Natl Acad Sci U S A 107:21140-5.

45. Willett JW, Kirby JR. 2012. Genetic and biochemical dissection of a HisKA domain identifies residues required exclusively for kinase and phosphatase activities. PLoS Genet 8:e1003084.

46. Saujet L, Monot M, Dupuy B, Soutourina O, Martin-Verstraete I. 2011. The key sigma factor of transition phase, SigH, controls sporulation, metabolism, and virulence factor expression in Clostridium difficile. J Bacteriol 193:3186-96.

47. Kint N, Janoir C, Monot M, Hoys S, Soutourina O, Dupuy B, Martin-Verstraete I. 2017. The alternative sigma factor sigma(B) plays a crucial role in adaptive strategies of Clostridium difficile during gut infection. Environ Microbiol 19:1933-1958.

48. Antunes A, Camiade E, Monot M, Courtois E, Barbut F, Sernova NV, Rodionov DA, Martin-Verstraete I, Dupuy B. 2012. Global transcriptional control by glucose and carbon regulator CcpA in Clostridium difficile. Nucleic Acids Res 40:10701-18.

49. Perego M, Hanstein C, Welsh KM, Djavakhishvili T, Glaser P, Hoch JA. 1994. Multiple protein-aspartate phosphatases provide a mechanism for the integration of diverse signals in the control of development in B. subtilis. Cell 79:1047-55.

50. Ohlsen KL, Grimsley JK, Hoch JA. 1994. Deactivation of the sporulation transcription factor Spo0A by the Spo0E protein phosphatase. Proc Natl Acad Sci U S A 91:1756-60.

51. Perego M. 2001. A new family of aspartyl phosphate phosphatases targeting the sporulation transcription factor Spo0A of Bacillus subtilis. Mol Microbiol 42:133-43.

52. Wang L, Grau R, Perego M, Hoch JA. 1997. A novel histidine kinase inhibitor regulating development in Bacillus subtilis. Genes Dev 11:2569-79.

53. Burkholder WF, Kurtser I, Grossman AD. 2001. Replication initiation proteins regulate a developmental checkpoint in Bacillus subtilis. Cell 104:269-79.

54. Mearls EB, Lynd LR. 2014. The identification of four histidine kinases that influence sporulation in Clostridium thermocellum. Anaerobe 28:109-19.

55. Obana N, Nakao R, Nagayama K, Nakamura K, Senpuku H, Nomura N. 2017. Immunoactive Clostridial Membrane Vesicle Production Is Regulated by a Sporulation Factor. Infect Immun 85.

56. Hussain HA, Roberts AP, Mullany P. 2005. Generation of an erythromycin-sensitive derivative of Clostridium difficile strain 630 (630Δerm) and demonstration that the conjugative transposon Tn916ΔE enters the genome of this strain at multiple sites. Journal of Medical Microbiology 54:137-141.

57. Thomas CM, Smith CA. 1987. Incompatibility group P plasmids: genetics, evolution, and use in genetic manipulation. Annual Reviews in Microbiology 41:77-101.

58. Yanisch-Perron C, Vieira J, Messing J. 1985. Improved M13 phage cloning vectors and host strains: nucleotide sequences of the M13mp18 and pUC19 vectors. Gene 33:103-19.

59. Lyras D, Rood JI. 1998. Conjugative Transfer of RP4-oriTShuttle Vectors fromEscherichia colitoClostridium perfringens. Plasmid 39:160-164.

60. McKee RW, Mangalea MR, Purcell EB, Borchardt EK, Tamayo R. 2013. The second messenger cyclic Di-GMP regulates Clostridium difficile toxin production by controlling expression of sigD. J Bacteriol 195:5174-85.

