## Supplemental Materials for "Three orphan histidine kinases inhibit *Clostridioides difficile* sporulation"

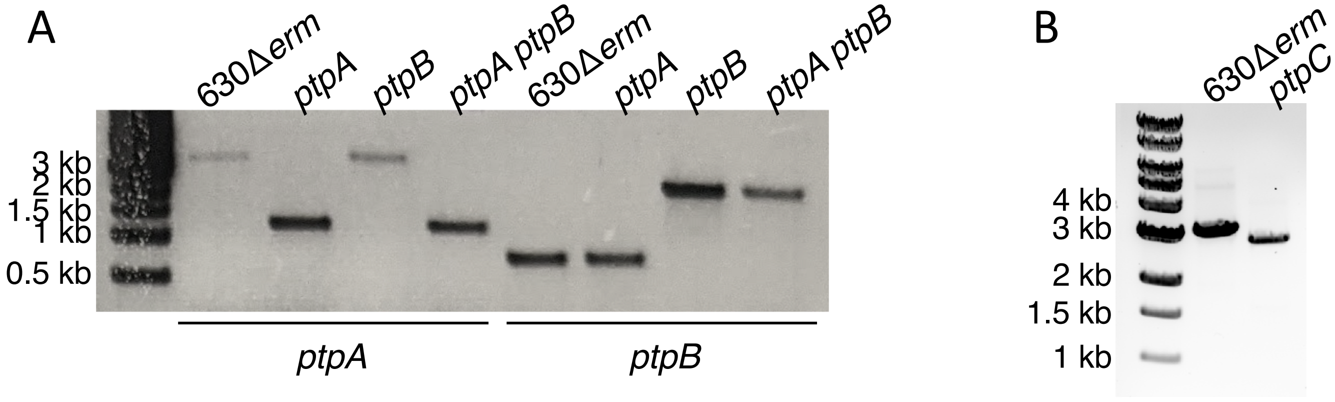


**Figure S1. PCR verification of *ptpA* (*CD1492*), *ptpB* (*CD2492*), *ptpA ptpB* (*CD1492 CD2492*) *and ptpC* (*CD1579*) mutants.** (**A**) PCR confirmation of *ptpA* deletion and *ptpB* disruption in 630Δ*erm*, *ptpA* (MC674), *ptpB* (MC788), and *ptpA ptpB* (MC802). (**B**) PCR confirmation of *ptpC* allelic replacement with *ermB* in 630Δ*erm* and *ptpC* (MC1646). Expected PCR products' sizes are: 4139 bp for the WT *ptpA* allele, 1400 bp for the *ptpA* deletion (primers oMC914/937); 811 bp for the WT *ptpB* allele, ~2800 bp for the *ptpB*::*erm* allele (primers oMC309/338); 3254 bp for the WT *ptpC* allele and 2921 bp for the *ptpC* allelic replacement (primers oMC2065/2066).

**Supplemental Figure 2. DNA cloning and vector details.**

pMC330: A construct encoding the group II intron from pCE240 to target insertion into *CD2492* at nucleotide 318s was amplified using primers oMC317, EBS1d_CD1A, oMC319 and EBSu and TA cloned into pCR2.1.

pMC333: The CD2492-specific group II intron from pMC330 was subcloned into pCE240 as *Bsr*GI/*Hind*III.

pMC336: The CD2492-specific group II intron from pMC333 was subcloned into pMC123 as *Sph*I/*Sfo*I.

pMC658: A 3011 bp fragment containing *CD2492* driven by its native promoter (194 bp) was amplified using primers oMC1481/oMC1482 and cloned into pMC123 using *Bam*HI/*Sph*I.

pMC673: A 3036 bp fragment containing *CD1492* driven by its native promoter (215 bp) expressed from its native promoter in pMC123 using primers oMC1537/1538 and BamHI/SphI

pMC681: A site-directed *CD2492-H664A* allele was synthesized and cloned into pUC19 by Genscript (Piscataway, NJ).

pMC683: A 3011 bp fragment containing the *CD2492-H664A* allele from pMC681 was subcloned into pMC123 as *Bam*HI/*Sph*I.

pMC707: A 2207 bp fragment containing *CD1579* driven by its native promoter was amplified with primers oMC1603/oMC1604 and cloned into pMC123 as *Bam*HI/*Sph*I.

pMC731: A 3036 bp fragment containing *CD1492* driven by its native promoter was amplified with primers oMC1749/oMC1750 and Gibson assembled into pMC658 as *Bam*HI/*Eco*RI.

pMC919: The ~740 bp upstream homology arm and ~500 bp downstream homology arms of *CD1579* were amplified with primers oMC1997/2139 and 0MC1998/2142 respectively. The *ermB* cassette from pJIR1457 was amplified using primers oMC2140/2141, and all three PCR fragments were Gibson assembled into pMSR as *Pme*I.

pMC982: The ~1950 bp and 347 bp fragments containing *CD1579* and introducing the H372A site-directed mutation were amplified using primers oMC1603/2499 and oMC1604/2498 and cloned into pMC123 as *Bam*HI/*Sph*I.

pMC1000: A site-directed CD1492-H668A was synthesized and cloned into pMC123 by Genscript (Piscataway, NJ).

pMC1062: An 140 bp fragment containing the sgRNA-*neg* sequence was amplified using primers 4084 and 4238 and cloned into pIA33 as *Msc*I/*Not*I.

pMC1064: An 140 bp fragment containing *sgRNA-CD1949-3* was amplified with primers oMC2787 and 4084 and cloned into pIA33 as *Msc*I/*Not*I.

pMC1095: An 140 bp fragment containing *sgRNA-CD1949-1* was amplified with primers oMC2785 and 4084 and cloned into pIA33 as *Msc*I/*Not*I.
